# Evaluation Of SYBR Green Real Time PCR For Detecting SARS-CoV-2 From Clinical Samples

**DOI:** 10.1101/2020.05.13.093609

**Authors:** Álvaro Fajardo, Marianoel Pereira-Gómez, Natalia Echeverría, Fernando López-Tort, Paula Perbolianachis, Fabián Aldunate, Pilar Moreno, Gonzalo Moratorio

**Affiliations:** Laboratorio de Virología Molecular, Centro de Investigaciones Nucleares, Facultad de Ciencias, Universidad de la República, Montevideo, Uruguay; Laboratorio de Evolución Experimental de Virus, Institut Pasteur, Montevideo, Uruguay; Laboratorio de Virología Molecular, Sede Salto, Centro Universitario Regional Litoral Norte, Universidad de la República, Salto, Uruguay

**Keywords:** SARS-CoV-2, molecular detection, SYBR Green, RT-qPCR

## Abstract

The pandemic caused by SARS-CoV-2 has triggered an extraordinary collapse of healthcare systems and hundred thousand of deaths worldwide. Following the declaration of the outbreak as a Public Health Emergency of International Concern by the World Health Organization (WHO) on January 30^th^, 2020, it has become imperative to develop diagnostic tools to reliably detect the virus in infected patients. Several methods based on real time reverse transcription polymerase chain reaction (RT-qPCR) for the detection of SARS-CoV-2 genomic RNA have been developed. In addition, these methods have been recommended by the WHO for laboratory diagnosis. Since all these protocols are based on the use of fluorogenic probes and one-step reagents (cDNA synthesis followed by PCR amplification in the same tube), these techniques can be difficult to perform given the limited supply of reagents in low and middle income countries. In the interest of economy, time and availability of chemicals and consumables, the SYBR Green-based detection was implemented to establish a convenient assay. Therefore, we adapted one of WHO recommended Taqman-based one-step real time PCR protocols (from the University of Hong Kong) to SYBR Green. Our results suggest that SYBR-Green detection represents a reliable cost-effective alternative to increase the testing capacity.

## INTRODUCTION

Ever since SARS-CoV-2 was identified as the etiological agent of a novel disease, COVID-19, at the beginning of the current year (Gorbalenya et al. 2020; Zhu et al. 2020a), the World Health Organization (WHO) has been following up on its spread (World Health Organization (WHO) 2020a). In addition, most of the scientific work has been mainly focused on three areas: i) the characterization of this virus and the disease that it caused; ii) the rapid developing of diagnostic methods; and iii) the patient treatments (Dennis Lo and Chiu 2020).

The rapid spreading of SARS-CoV-2 highlights the need for an effective surveillance method to be widely used in different laboratory settings (Thompson 2020). This fact has prompted the development of a wide variety of molecular diagnostic methods based on the detection of viral genomic RNA. The vast majority rely on reverse transcription real time PCR (RT-qPCR), due to its high sensitivity and specificity (Chu et al. 2020; Corman et al. 2020; Huang et al. 2020; World Health Organization (WHO) 2020b; Zhu et al. 2020a). This technique, either as a one-step or a two-step protocol, has accelerated PCR laboratory procedures and has had the strongest impact on virology as it is being applied for detection, quantification, differentiation and genotyping of animal and human viruses (Bankowski and Anderson 2004; Kaltenboeck and Wang 2005). Furthermore, it is regarded as a gold standard for analysis and quantification of pathogenic RNA viruses in clinical diagnosis (Espy et al. 2006). In particular, for the molecular diagnosis of COVID-19 the WHO website recommends few One-step RT-qPCR detection protocols that have been developed in different countries (World Health Organization (WHO) 2020b). Since all these protocols are based on the use of fluorogenic probes and one-step reagents (cDNA synthesis followed by PCR amplification in the same tube), these techniques are limited to the use of more specific reagents and can be quite expensive. Moreover, these protocols involve the amplification of more than one gene, which implies different probes and fluorescent channels.

Therefore, several researchers have attempted to develop alternative SARS-CoV-2 detection methods that might be faster or cheaper to implement, such as loop-mediated isothermal amplification (LAMP) (Jiang et al. 2020; Park et al. 2020; Yang et al. 2020; Zhang et al. 2020; Zhu et al. 2020b), droplet digital PCR (ddPCR) (Dong et al. 2020; Suo et al. 2020), multiplex PCR (Li et al. 2020) or even protocols based on CRISPR-Cas12 (Curti et al. 2020). Furthermore, considering the shortage in the supply of RNA extraction kits, others have evaluated alternative nucleic acids extraction methods (Bruce et al. 2020; Ladha et al. 2020; Zhao et al. 2020). Despite all these approaches, there is still much to be done to generate strategies that might be helpful for different laboratory settings.

Quantitative PCR (qPCR) is a molecular technique widely used when detection and/or quantification of a specific DNA target is needed. qPCR is based on fluorescence to measure the amount of a DNA target present at each cycle of amplification during the PCR. The most common ways of generating a fluorescent signal are by use of specific hydrolysis probes (i.e. TaqMan® probes), or a double-stranded DNA binding dye (i.e SYBR® Green). SYBR-Green-based detection method presents several advantages over Taqman chemistry ones, as being cheaper and not requiring the synthesis of specific probes.

This technique has already been proposed and used for laboratory testing of different pathogens, including viruses (Espy et al. 2006; Fernández et al. 2006; Gomes-Ruiz et al. 2006; Kumar et al. 2012), bacteria (Kositanont et al. 2007; Keerthirathne et al. 2016) and unicellular protozoan parasites (Espy et al. 2006; Haanshuus et al. 2019), among others. For SARS-CoV-2 detection, some preliminary reports have attempted to assess the sensitivity and predictive value of the different sets of primers and probes available (either commercially or *in-house* developed) (Barra et al. 2020; Casto et al. 2020; Jung et al. 2020), but so far, no comparison has been made between the different real time chemistries for this emerging virus.

The aim of the study was to set up an alternative molecular protocol to detect SARS-CoV-2 from clinical samples, without the need of TaqMan probes or post-PCR steps (i.e. gel electrophoresis), which can be implemented in case of difficulties to get specific reagents or kits because of the current pandemic situation. Here we showed that Taqman-based one-step real time PCR protocol recommended by the WHO (Chu et al. 2020; Poon et al. 2020) can be successfully adapted and alternatively used with SYBR Green-based two-step qPCR.. Besides, performing a comparison of the different molecular techniques by employing dilutions of control vectors and RNA standards for quantification, we tested our assay with 8 clinical samples collected from confirmed COVID-19 cases and one negative patient. Overall, our results showed that both approaches were able to detect SARS-CoV-2 from clinical samples.

## MATERIALS AND METHODS

### Positive controls, clinical samples and ethical considerations

Positive controls were kindly provided by Dr. Leo Poon from the University of Hong Kong. Positive controls contain a region of ORF1b-nsp14 or N targets of SARS-CoV Urban strain cloned into a standard plasmid. Residual de-identified nasopharyngeal samples were remitted to the Institut Pasteur Montevideo, that has been validated by the Ministry of Health of Uruguay as an approved center providing diagnostic testing for COVID-19.

### SARS-CoV-2 One Step RT-qPCR protocol with fluorogenic probes

The one-step RT-qPCR protocol evaluated in this study corresponded to the one developed by the University of Hong Kong (Chu et al. 2020; Poon et al. 2020), with modifications, which consists of two monoplex real-time RT-PCR assays targeting the ORF1b-nsp14 and N gene regions of SARS-CoV-2 (Supplementary Table 1). Concentrations used were lowered to avoid non-specific amplification (data not shown). Briefly, a 20μL monoplex reaction contained 5μL of 4x TaqMan Fast Virus Master Mix (Thermo Fisher), 0.6µL of each primer (0.3μM final concentration each), 0.2μL of the probe (0.1μM final concentration) and 4μL of RNA. These monoplexes were performed for both N and ORF1b-nsp14 regions. Thermal cycling was run on a Step-One Plus RT-PCR thermal cycler (Applied Biosystems) with the following cycle parameters: 50°C for 5min for reverse transcription, inactivation of reverse transcriptase at 95°C for 20s and then 40 cycles of 95°C for 5s and 60°C for 30s. The expected amplicon sizes of ORF1b-nsp14 and N are 132bp and 110bp, respectively. This protocol was carried out with serial dilutions of plasmids containing N and ORF1b-nsp14 genes (kindly provided by Dr. Leo Poon from the School of Public Health, University of Hong Kong), RNA standards of the same targets constructed in our laboratory, and later validated with RNA samples of COVID-19 cases. A non-template control (nuclease-free water) was included in every one-step RT-qPCR run. We manually set the threshold value to 0.015 in all assays to determine the threshold cycle (C_t_). A test run for the amplification of the controls was done to select the appropriate dilution to use for the amplification of the clinical samples, which were, in turn, run in duplicates.

**Table 1.** Ct values and melting temperatures (Tm) of the amplified control dilutions according to the probe-based RT-qPCR protocol versus the SYBR Green-based qPCR protocol developed in this work.

| Target | Copy number | Probe-based | SYBR Green-based |  |
| --- | --- | --- | --- | --- |
|  |  | Ct (Threshold 0.015) | Ct (Threshold 0.2) | T<sub>m</sub> (°C) |
| ORF1b-nsp14 | 10<sup>4</sup> | 35.79 | 31.08 | 81.55 |
|  | 10<sup>5</sup> | 30.78 | 26.11 | 81.40 |
|  | 10<sup>6</sup> | 26.93 | 22.54 | 81.40 |
|  | 10<sup>7</sup> | 23.38 | 18.40 | 81.40 |
|  | C- | 40.00 | 40.00 |  |
| N | 10<sup>4</sup> | 37.07 | 31.27 | 81.70 |
|  | 10<sup>5</sup> | 31.52 | 26.03 | 81.70 |
|  | 10<sup>6</sup> | 28.19 | 21.93 | 81.70 |
|  | 10<sup>7</sup> | 23.47 | 17.61 | 81.70 |
|  | C- | 40.00 | 37.76 | 71.57* |

C -: non-template-control
\* Non-specific amplification

### SARS-CoV-2 qPCR protocol with SYBR Green

First, complementary cDNA of SARS-CoV-2 clinical samples was generated using SuperScript II Reverse Transcriptase (Invitrogen), random primers and 10μL of RNA, according to the manufacturer’s instructions.

qPCR reactions were carried out using a Step-One Plus RT-PCR thermal cycler (Applied Biosystems), Luna Universal qPCR Master Mix (New England Biolabs), following manufacturer’s instructions, and the same primers previously used in the One Step RT-qPCR Taqman protocol. Each 20μl reaction contained 10μL of 2x Master Mix (NEB), 0.5µL of each primer (0.25μM final concentration each) and 4μL of cDNA. Again, non-template control (nuclease-free water) was included in every qPCR run as a negative control. In this case, we set the threshold value to 0.2 in all assays to determine the C_t_. As with the probe-based protocol, a test run for the amplification of the control plasmids was done to select the appropriate dilution to use for the amplification of the clinical samples, which were, in turn, run in duplicates. The cycling conditions were: initial denaturation at 95°C for 1min, 40 PCR cycles of 95°C for 15s and 60°C for 30s, followed by a melting curve ranging from 60°C to 95°C (acquiring fluorescence data every 0.3°C). With the aim of verifying specific amplification, in addition to the melting curve step during the run, we also confirmed the amplicon sizes by 2% agarose gel electrophoresis.

### Construction of RNA for quantification standards

A fragment of 132 and 110 bp containing the ORF1b-nsp14 and N targets, respectively, were cloned into pCR(tm)2.1-TOPO® using the TOPO® TA Cloning® Kit (Invitrogen) following manufacturer’s instructions and transformed in NEB® 5-alpha Competent *E. coli* (High Efficiency) by the heat shock method (42°C, 30 s). Plasmids were isolated using PureLink Quick Plasmid Miniprep Kit (Invitrogen) and quantified by spectrophotometric analysis (Biophotometer, Eppendorf). Then, 1µg of each plasmid was linearized with *Spe*I and *in vitro* transcribed with T7 RNA Polymerase (Thermo Fisher) following the manufacturer’s instructions. *In vitro* transcribed RNA was treated with DNase and purified with TURBO DNA-free(tm) Kit (Thermo Fisher). RNA purified was checked for size and integrity by gel electrophoresis. The number of copies/µL was calculated as: (N_A_ x C)/ MW, where, N_A_ is the Avogadro constant expressed in mol^−1^, C is the concentration expressed in g/µL, and MW is the molecular weight expressed in g/mol.

### Determination of the sensitivity of the assays by standard curves

A stock containing around 2 × 10^13^ copies/µL (for both ORF1b-nsp14 and N) was used for standard curve and sensitivity determination of the qPCR assays. The standard curve and sensitivity were determined by 10-fold serial dilutions. In the case of the one-step probe-based qPCR assays, 4µL of RNA was directly added to the mix and run, in triplicates, as mentioned above. For the two-step SYBR Green-based qPCR 4µL of the same 10-fold serial dilutions of the *in vitro* transcribed RNA for each target were retrotranscribed and then 4µL of the cDNA was used as template for SYBR Green qPCR. Each cDNA was run in duplicates. Standard curves were represented as C_t_ vs log copy number/reaction. The lower limit of detection was defined as the lowest copy number of target/qPCR, taking account for dilution, which amplified reliably.

### Molecular cloning of N amplicons from clinical samples and Sanger sequencing

PCR products generated by the qPCR protocol with SYBR Green contain dA overhangs at the 3’ ends. Therefore, the fresh PCR products of the N target from samples 1, 3, 4 and 6 were directly cloned into pCR(tm)2.1-TOPO® using the TOPO® TA Cloning® Kit (Invitrogen) following manufacturer’s instructions. Next, cloning reactions were transformed in NEB® 5-alpha Competent *E. coli* (High Efficiency) by the heat shock method (42°C, 30 s), plated in LB medium containing 50 µg/mL ampicillin (Amp), 40µL Xgal (40 mg/mL), 10 µL IPTG (100 mM) and incubated at 37°C overnight. Three individual white colonies for each cloning reaction were isolated and overnight cultured in LB containing 50 µg/mL ampicillin. Plasmids were isolated using PureLink Quick Plasmid Miniprep Kit (Invitrogen) and Sanger sequenced with the universal primers M13Forward and M13Reverse.

### Sequences analysis

Ab1 files from Sanger sequencing were analyzed using the Staden package v1.7.0 (http://staden.sourceforge.net). MEGAX (http://www.megasoftware.net) was used to perform sequence analysis.

### *In silico* estimation of the DNA Melting Temperature

GC content and DNA melting temperature of ORF1b-nsp14 and N targets from SARS Urbani isolate (MK062184) and SARS-CoV-2 (MT358402) were estimated using the DNA Melting Temperature (Tm) Calculator (available at http://www.endmemo.com/bio/tm.php). The rationale for setting the parameters was to emulate as much as possible SYBR-Green qPCR conditions used in this study. To do so, the salt and magnesium concentration were set at 50 mM and 1.5 mM, respectively, as it is indicated by the manufacturer. Initial DNA copy numbers were obtained using the C_t_ values empirically observed in the SYBR Green qPCR for the positive control and the clinical samples for both targets (showed in Table 2). Then, we interpolated them in their corresponding standard curve in order to estimate the initial number of copies in the qPCR reaction. After this, we calculated the number of target copy numbers after one PCR cycle and estimated the DNA concentration for each sequence expressed as nM using the DNA/RNA Copy Number Calculator from the http://www.endmemo.com/bio/dnacopynum.php website.

**Table 2.**
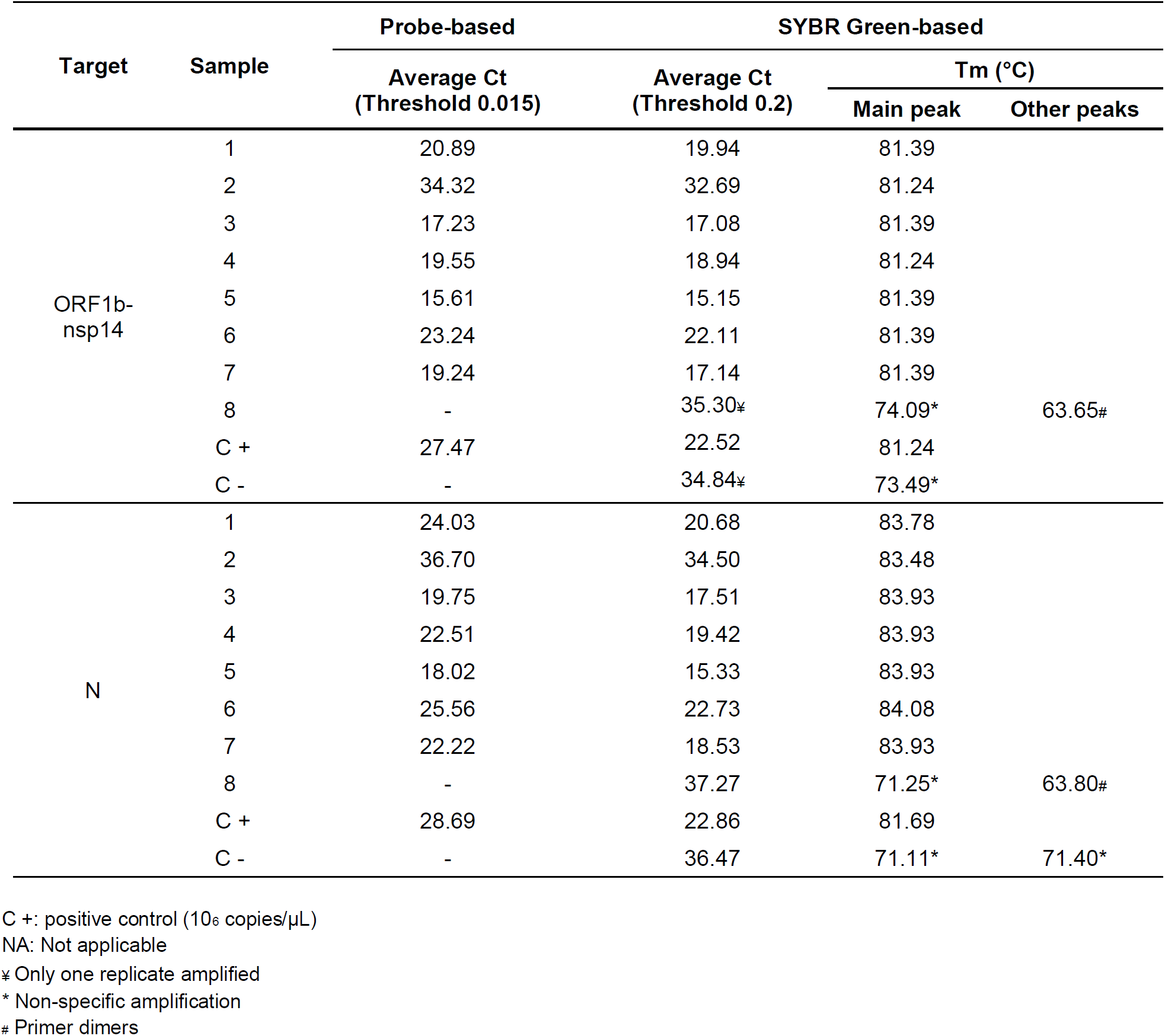
Average C_t_ values and melting temperatures (Tm) of the clinical samples (run in duplicates) according to the probe-based RT-qPCR protocol versus the SYBR Green-based method developed in this work.

## RESULTS

### Set up of qPCR protocols (SYBER Geen and Taqman chemistry) with DNA controls for ORF1b-nsp14 and N targets

In order to select an appropriate amount of control vector to use in the comparison between the two real time qPCR methods, we prepared plasmids dilutions (10^7^, 10^6^, 10^5^ and 10^4^ copies/μL) and assayed them following both protocols: the probe-based One Step RT-qPCR developed by the University of Hong Kong (Chu et al. 2020; Poon et al. 2020) and the *in-house* SYBR Green-based protocol adapted in this study. It is worth mentioning that previous results, of our laboratory, had indicated that a lower amount of primers and probes than initially suggested by Poon et al. (2020), rendered similar positive results, and diminished the amplification of primer dimers (data not shown). Real time PCR results, from SYBER and Taqman chemistries, of different dilutions of the control vectors for the targeted regions (ORF1b-nsp14 and N) are shown in Figure 1 (panels A, B, C and D). Since all dilutions amplified correctly and below a Ct of 37 (Fig. 1 and Table 1), we decided to use 10^6^ copy number/μL as a positive control for subsequent assays (for both ORF1b-nsp14 and N genes).

**Figure 1.**
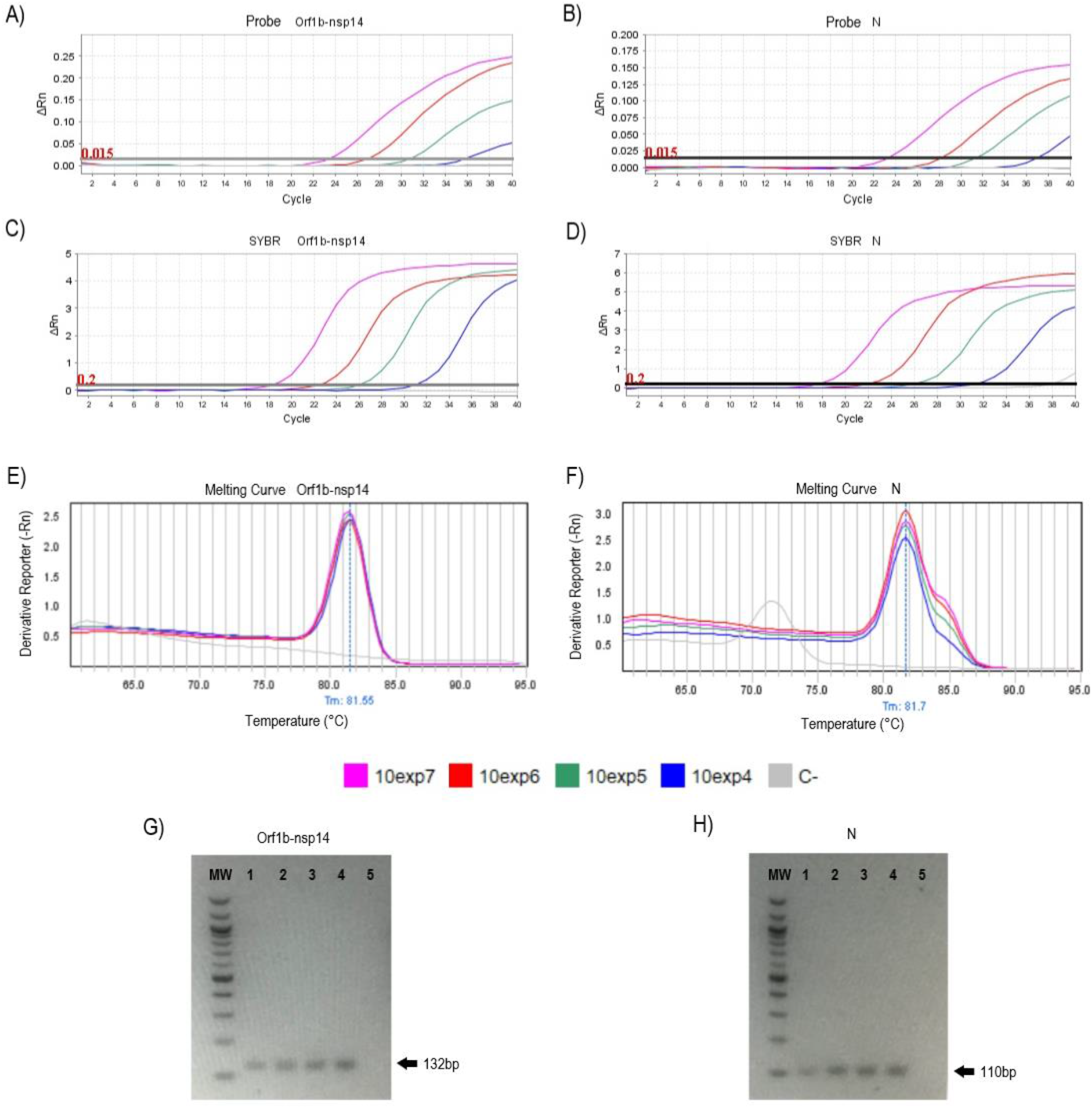
Real time PCR results, from SYBER and Taqman chemistries, of different dilutions of the control vectors for the targeted regions: ORF1b-nsp14 and N. (left and right panels, respectively). **A)** and **B)** show the amplification plots for the RT-qPCR protocol employing fluorogenic probes. **C)** and **D)** show the amplification plots for the qPCR protocol developed in this study employing SYBR Green as a nucleic acid dye. **E)** and **F)** show the melting curves for the products amplified with the SYBR Green-based qPCR protocol. Below these panels are the references for each of the dilutions assayed expressed in plasmid copies (C-: non-template control). **G)** and **H)** show agarose gel electrophoresis of PCR products amplified with the SYBR Green-based qPCR protocol. MW: 100bp DNA Molecular Weight (New England Biolabs); lanes 1 to 4: control dilutions (10^4^, 10^5^, 10^6^ and 10^7^ copies/μL, respectively); lane 5: non-template-control.

Analyzing the specificity of the SYBR Green-based qPCR method (Fig. 1, panels E to H) from ORF1b-nsp14, we verify the presenceof only one PCR product, corroborated by a unique melting peak (Tm=81.55°C) (Fig. 1E and Table 1). Agarose gel electrophoresis (Fig. 1G) allowed also the verification of the expected product size (132bp) with no amplification in the negative control. In the case of the SYBR Green-based qPCR method for N gene amplification we can observed (Fig. 1D and F, for all N dilutions, a very clear peak at Tm=81.70°C, together with a non-symmetric melting temperature peak slightly skewed to a higher temperature, which might suggest the presen ce of two PCR products. However, when we run the PCR products on an agarose gel only one product of the expected size (110bp) is observed (Fig 1H), demonstrating that the presence of a double peak was not indicative of non-specific amplification.. For the non-template-control we observed a slight amplification (Ct=37.76), although the melting curve evidenced a non-specific peak (Tm=71.57°C)

### Validation of both qPCR methods using clinical samples

To validate the SYBR Green qPCR protocol, we assayed a set of 8 RNA samples from COVID-19 cases with both qPCR methods and both genetic regions. These samples were beforehand determined as SARS-CoV-2 positive (samples 1 to 7) and negative (sample 8), employing the diagnostic kit provided by the Panamerican Health Organization (Berlin Protocol). The results obtained, with both qPCR assays chemistries and genetic regions, were in agreement with the data previously gatheredfor these samples (Fig. 2 and 3)..

**Figure 2.**
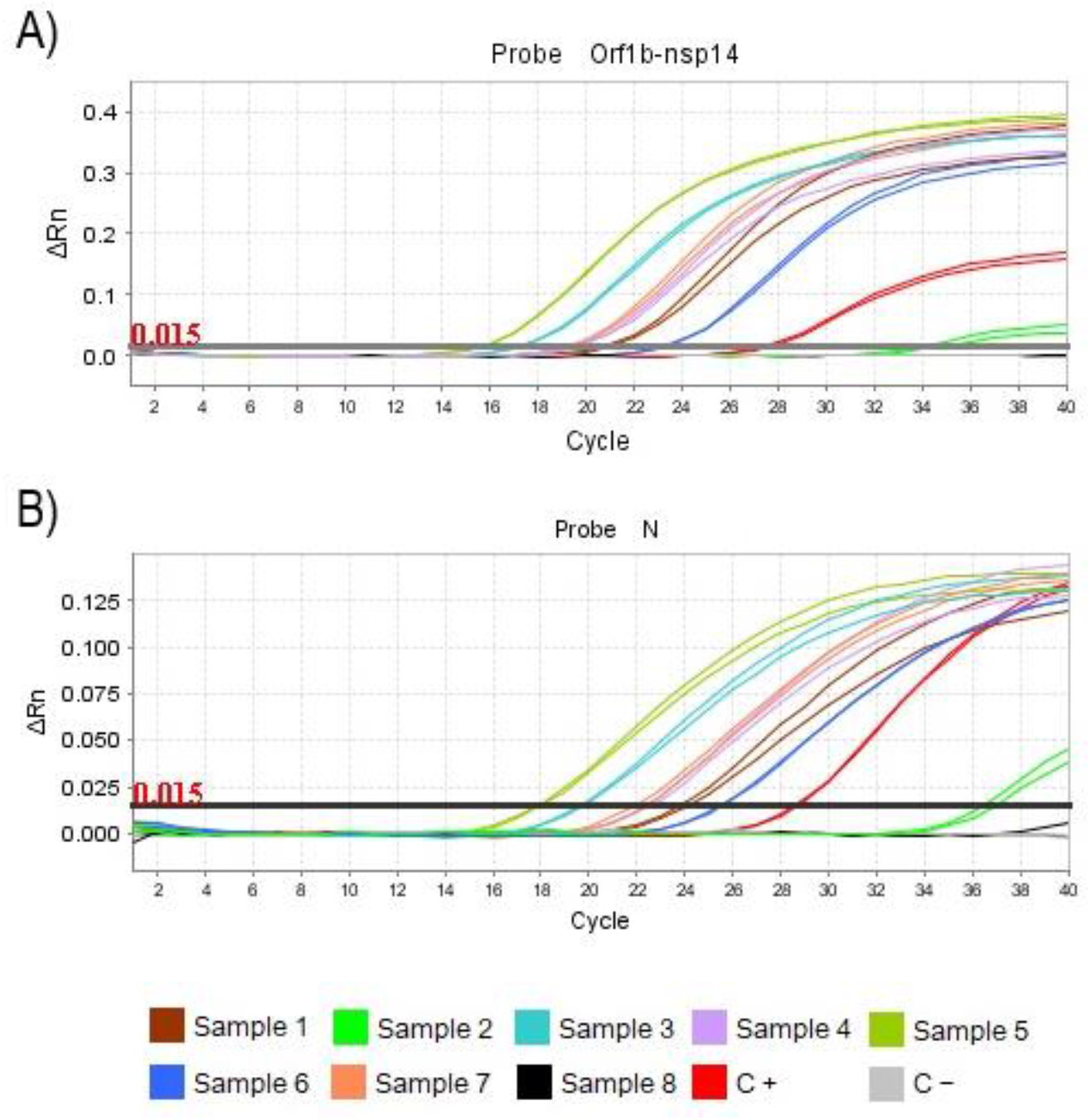
Amplification plots for the RT-qPCR protocol employing Taqman probes to detect SARS-CoV-2 in clinical samples. Adapted method previously described by Poon et al. 2020. As a positive control (C+), 10^6^ copies/μL of each vector were used. DEPC water was used as non-template-control (C-),. Each sample was assayed in duplicate. **A)** Detection of ORF1b-nsp14 region. **B)** Detection of N region. The references for both panels are indicated below panel B.

**Figure 3.**
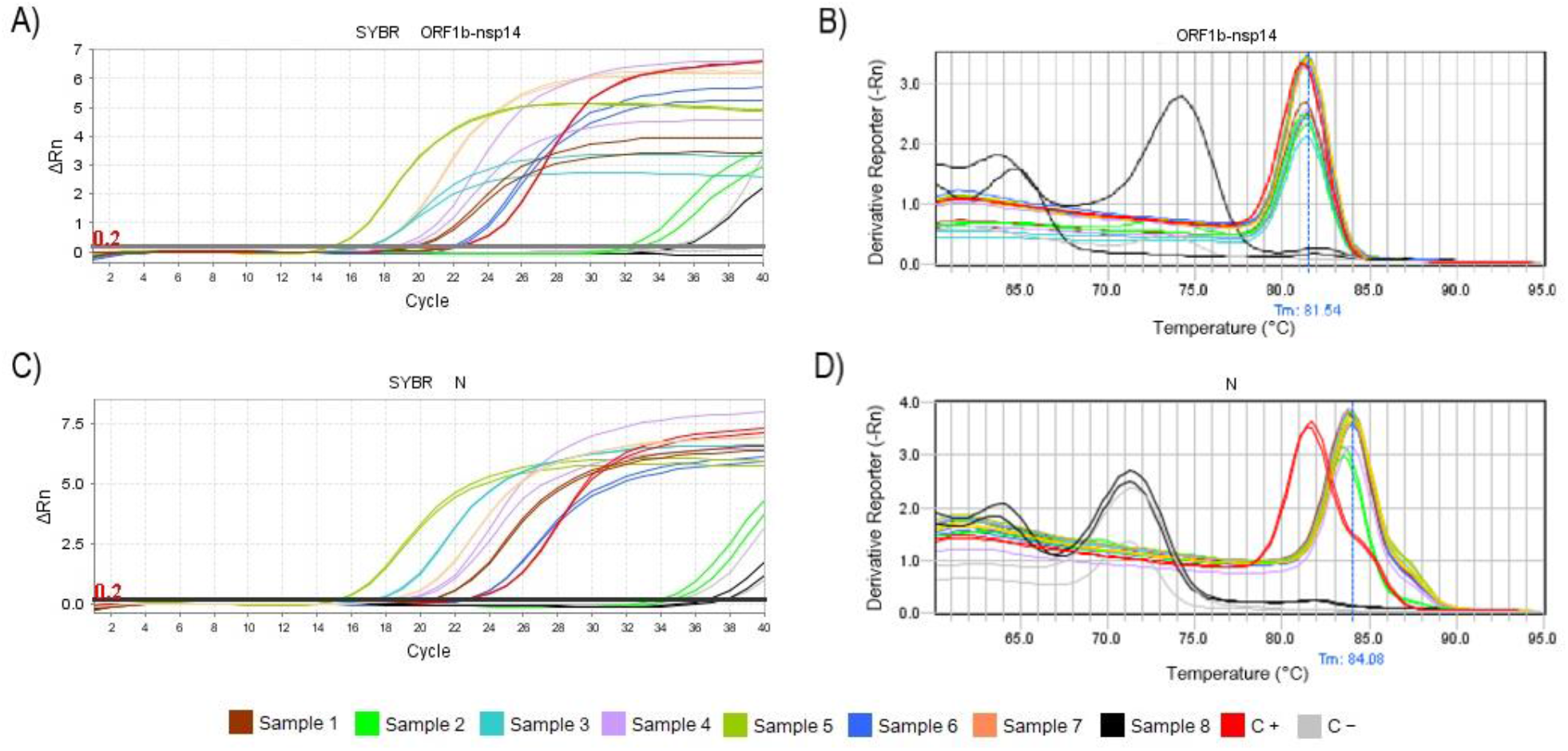
qPCR results of the protocol employing SYBR Green to detect SARS-CoV-2 in clinical samples. As a positive control (C+), 10^6^ copies/μL of each vector were used., DEPC water was used as a non-template-control (C-). Each sample was assayed in duplicate. **A)** Amplification Plot and **B)** Melting curve for the detection of ORF1b-nsp14 region, respectively. **C)** Amplification Plot and **D)** Melting curve for the detection of N region, respectively. The references for all panels are indicated at the bottom of the figure.

The amplification data for the SYBR Green-based qPCR protocol showed that the ORF1b-nsp14 region was correctly amplified for all SARS-CoV-2 positive samples (1 to 7) (Fig. 3). This was verified by melting curve analysis in every case, in agreement with the positive control (Fig. 3B and Table 2). In addition, sample 2, which was suspected to have a low viral load, was correctly amplified with this protocol. As for the negative viral RNA sample (sample 8), even though it seems to amplify in very late cycles (Fig. 3A and Table 2), the melting curve analysis reveals that the amplification corresponds to primer dimers and/or non-specific products (Fig. 3B and Table 2).

As in the case of ORF1b-nsp14 amplification, the amplification of N region allowed the correct assignment of all positive and negative clinical samples. The non-template controls, as well as sample 8 (negative for SARS-CoV-2), showed delayed non-specific amplification (Fig. 3C and 3D, and Table 2), which in principle does not invalidate the results, because are in agreement with the results of the assay using different dilutions of the control vectors. In addition, the clinical samples showeda skewed peak similar to the one that was previously observed for the positive control. (Fig. 3D).Taking together, these results suggest the specific amplification of N target. Despite all SARS-CoV-2 positive samples amplify the same product, they exhibit a higher Tm than the positive control (83.86 ± 0.07°C vs 81.69°C, respectively), effect that itwas not observed for ORF1b-nsp14 target. This result can be explained as the positive control used in this study correspond to a SARS-CoV Urbani isolate (Genbank Accession number MK062184) which was used due to the unavailability of a SARS-CoV-2 positive control.

### Tm differences observed, in the N target amplification, between positive control and clinical samples are explained by differences in their GC content

The Tm of a DNA fragment depends on a variety of features such as its length, GC composition, sequence and concentration, among others. Given the results previously described, we hypothesized that the difference between the Tm of the positive control and the clinical samples assayed here was due to a higher GC content in the N target from Uruguayan patients. To test this, we first cloned 5 of 7 PCR product obtained from the clinical samples and sequenced one molecular clone for each (Figure 4). The results showed that all sequences were identical between them and a BLAST search showed that all cloned sequences had 100% identity with SARS-CoV-2, confirming that SYBR-Green based qPCR specific amplified viral RNA present in the clinical samples.

**Figure 4.**
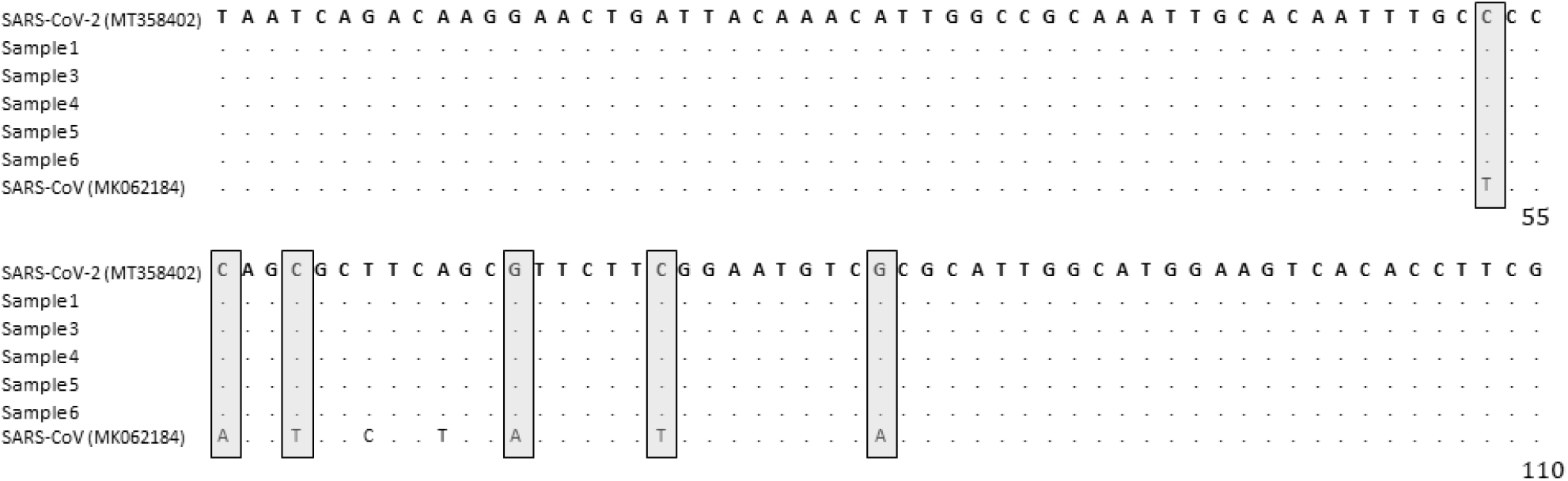
Sequences of molecular clones from the clinical samples. Sequences alignment of the N target amplicon for SARS-CoV-2, SARS-CoV (positive control) and molecular clones obtained from clinical samples used in this study. All clones fully matched with the SARS-CoV-2 sequence. Below is shown the sequence of the N target for the positive control. Boxes indicate nucleotide positions (6/110) which contribute to an increment in the GC content of the SARS-CoV-2 sequences, compared to the sequence of SARS-CoV. Accession numbers are indicated between brackets.

We then calculated the GC content as well as the Tm was *in silico* estimated for the amplicon sequences of the N target from clinical samples. Comparisons were made taking as reference the the N target of the SARS-CoV Urbani strain, which was used as positive control (Table 3). Given the *in silico* Tm estimates, the N amplicons obtained for SARS-CoV-2 samples should have a higher Tm (around 2°C) than that of SARS-CoV Urbani strain, which confirms that the Tm differences observed for N amplicon derive from their different GC content. The magnitude of the Tm gain correlated positively with the GC% (Pearson, r = 0.843, P < 0.001). Therefore, we conclude that the observed differences on the Tm of N targets from clinical samples were due to differences in their GC content.

**Table 3.** GC content and differences between Tm empirically observed and *in silico* estimated for N amplicons of SARS-CoV (positive control) and SARS-CoV-2 (clinical samples) in SYBR Green-based qPCR assays. Observed values for positive control and clinical samples were taken from Table 2. Observed values for clinical samples were averaged.

| Target | Sample | GC (%) | Ct observed | $T_m$ observed (°C) | $T_m$ estimated (°C) |
| --- | --- | --- | --- | --- | --- |
| N | C+ | 44.55 | 22.86 | 81.69 | 81.12 |
| | Clinical samples | 49.09 | $21.24 \pm 2.21$ | $83.86 \pm 0.07$ | 83.10 |

### Limit of quantitation of Taqman probe and SYBR-Green based qPCRs

Finally, to determine the limit of quantitation of both SARS-CoV-2 detection qPCR approaches, serial dilutions of an RNA standard for each target were performed (Figure 5). As expected, the C_t_ of each reaction increased along with the lower number of target copies/reaction. The C_t_ values showed an inverse linear relationship with the log value of the RNA concentrations with a very high correlation (R^2^ > 0.99 in all cases). The results showed that the limit of quantitation for ORF1b-nsp14 and N targets were equal to 10^3^ copies/reaction for probe-based qPCR (Figure 5A and 5C) and 20×10^3^ copies/reaction (Figure 5B and 5D) for SYBR-Green qPCR assays.

**Figure 5.**
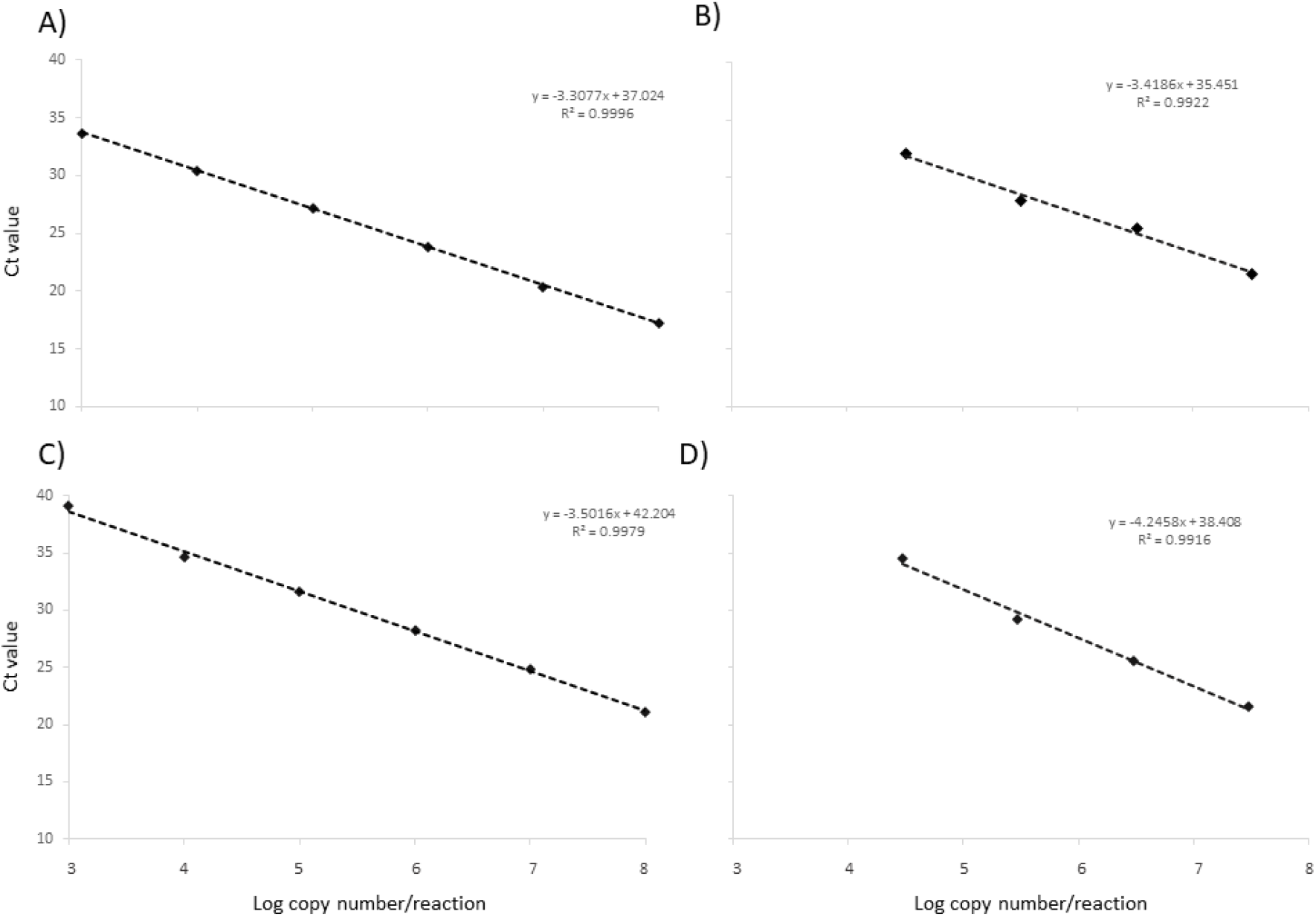
Standard curves of Taqman and SYBR Green based qPCR for targets ORF1b-nsp14 and N. Serially diluted RNA containing ORF1b-nsp14 (A, B) or N (C, D) targets were amplified and analyzed in both one-step (A, C) and two-step (B, D) qPCR protocols. The threshold cycle (C_t_) mean values were plotted against copy number of RNA standards/reaction. The coefficient of determination (R^2^) and the lineal regression curve (y) were determined. Each dilution was performed in triplicates (Taqman assays) or duplicates (SYBR Green assays).

## DISCUSSION

The qPCR technique is widely used in clinical virology diagnostic laboratories because of its high sensitivity, specificity, reproducibility and no need of post PCR steps (Josko 2010). Additionally, qPCR allows for quantification because during the amplification of the target, it reaches a threshold level that correlates with the amount of initial target sequence (Valasek and Repa 2005). SYBR-Green based qPCR has relatively low cost benefit, whereas Taqman-based qPCR are more expensive. In addition, the specificity of the qPCR is mainly provided by the use of specific primers, although Taqman probes increase the specificity because only sequence-specific amplifications are measured (Tajadini et al. 2014).

Our results with SYBR Green chemistry were consistent with the initial probe-based protocol designed by Poon et al. (2020) showing sensitivity to specifically detect SARS-CoV-2. It is worth noting that the original protocol (Chu et al. 2020; Poon et al. 2020) suggested the use of the N target for screening analyses, whereas the amplification of ORF1b-nsp14 was indicated as a confirmatory assay. ORF-nsp14 encodes for a very conserved exoribonuclease present in all known coronaviruses which is involved in replication fidelity (Eckerle et al. 2007, 2010). The N gene encodes for the structural nucleoprotein, which is more exposed to the recognition of the host immune system and therefore could be more prone to change than ORF-nsp14 (Woo et al. 2010). Importantly, if mutations occur within the probe-binding site, they would prevent the annealing of the probe and its subsequent detection. Although coronaviruses are between the RNA viruses with lower mutation rates (Sanjuan et al. 2010), it would be possible that new mutations impact negatively on the probe detection. Therefore, counting on an alternative detection method such as SYBR Green-based two-step qPCR, that only requires two conserved regions for primer binding instead of three (for hybridization probe), it might become useful. In this context, our results with SYBR Green chemistry may provide a simpler and cheaper alternative for SARS-CoV-2 detection.

Here we reported a lower limit of detection of the Taqman probe-based approach compared to the SYBR-Green based qPCR. In addition to the decrease in specificity due to the lack of use of a probe, SYBR Green-based qPCR approach needs a previous step of cDNA synthesis. This extra step represents a possible source of contamination that can affect the results. In order to increase the specificity of the SYBR Green-based qPCR assayed here, we could evaluate the use of specific primer instead of random hexamers during the retrotranscription step. Another disadvantage of the SYBR Green vs probe-based qPCR is that any non-specific product including primer-dimer can lead to false positive results. For this reason, the melting curve analysis must be performed to confirm that only specific amplification was obtained.

Although multiplexed qPCR is more frequently developed for Taqman technology, our results suggest that a multiplexed SYBR Green-based qPCR could be developed for SARS-CoV-2 detection. The difference of the GC% content among the targets Orf-nsp14 and N which produce a Tm difference of 2°C, seems to be enough for simultaneous detection of both targets in the same tube. In this case, after melting curve analysis two specific double peaks should be observed.

Altogether, both SYBR Green-based qPCR and Taqman probe-based qPCR assays for detecting SARS-CoV-2 were set up in our laboratory conditions and their consistencies, as well as their advantages and disadvantages, were analyzed. This work could help to increase the testing capacity of some places in the world with limited access to Taqman specific reagents, given the current lockdown of many countries.

## ACKNOWLEDGEMENTS

We thank Dr. Leo Poon and his group (School of Public Health, The University of Hong Kong) for kindly providing us with the control vectors for ORF1b-nsp14 and N regions of SARS-CoV used in this work.

## AUTHORS’ CONTRIBUTIONS

PM, GM conceptualized the study design; AF, FLT, FA, PP, MP-G performed the laboratory tests; NE and MP-G plotted the figures; AF, FLT, FA, PP, NE, MP-G, PM, GM analyzed the data and interpreted the results; NE, MP-G, PM and GM wrote the manuscript. All authors read and approved the final report.

---

**Supplementary Table 1.**
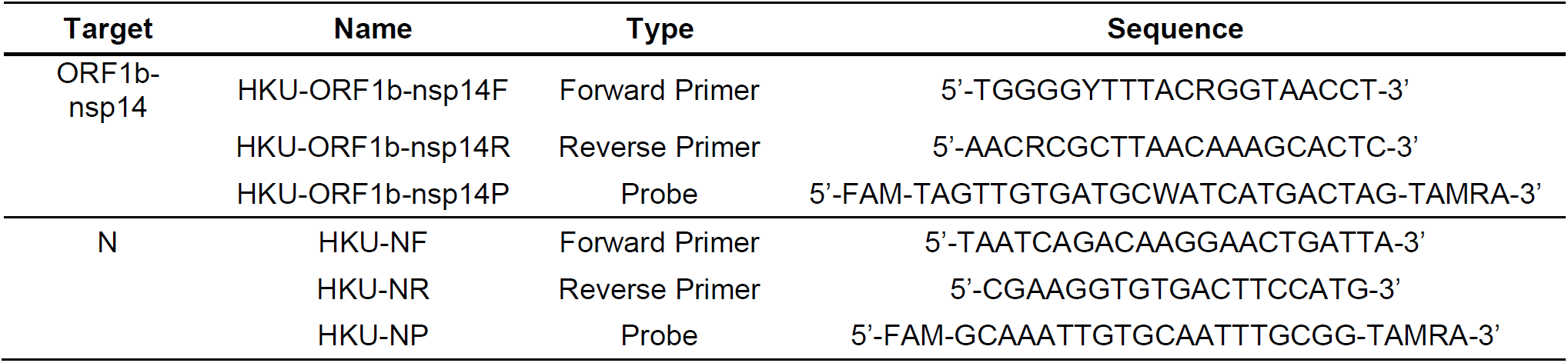
Information of primers and probes tested in this study from the University of Hong Kong protocol (Poon et al. 2020).

